# Interplay of primary bovine lymphocytes and *Mycobacterium avium* subsp. *paratuberculosis* shows distinctly different proteome changes and immune pathways in host-pathogen interaction

**DOI:** 10.1101/578187

**Authors:** Kristina J.H. Kleinwort, Stefanie M. Hauck, Roxane L. Degroote, Armin M. Scholz, Christina Hölzel, Erwin P. Märtlbauer, Cornelia A. Deeg

**Author notes:** Corresponding author: Cornelia A. Deeg, Chair for Animal Physiology, Department of Veterinary Sciences, LMU Munich, Veterinärstr. 13, D-80539 Munich.

## Abstract

Mycobacterium avium subsp. paratuberculosis (MAP) is a pathogen causing paratuberculosis in cattle and small ruminants. During the long asymptomatic subclinical stage, high numbers of MAP are excreted and can be transmitted to food, where they survive many of the standard techniques of food decontamination. If these MAP are harmful to the consumers is currently under debate. In general, there is a lack of information regarding interaction of the hosts immune system with MAP.

In this study, we tested the interaction of peripheral blood lymphocytes (PBL) from cattle with MAP in their exoproteomes/secretomes. Because in other mycobacterial infections, the immune phenotype correlates with susceptibility, we additionally tested the interaction of MAP with recently detected immune deviant cows.

In PBL, different biological pathways were enhanced in response to MAP dependent on the immune phenotype of the host. PBL of control cows activated members of cell activation and chemotaxis of leukocytes pathway as well as IL-12 mediated signaling. In contrast, in ID cows CNOT1 was detected as highly abundant protein, pointing to a different immune response, which could be favorable for MAP. Additionally, MAP reacted different to the hosts. Their exoproteomes differed in either GroEL1 or DnaK abundance, depending on the interacting immune response.

These findings point to an interdependent, tightly regulated response of MAP and the immune system.

## 1. Introduction

*Mycobacterium avium* subsp. *paratuberculosis* (MAP) is a critical pathogen for cattle and small ruminants, causing paratuberculosis with decreased milk production and, in some animals, excessive loss of weight [1]. Paratuberculosis, also referred to as Johne’s disease, is world-wide endemic; no country or region has been found to be free of this disease [2]. Affected ruminants go through a long asymptomatic subclinical phase in which infection cannot reliably be detected by standard diagnostic tests [3, 4]. These subclinical infected animals can already shed MAP, thereby contaminating dairy products or meat [5]. Diseased animals were shown to shed high numbers of MAP [6, 7]. Since viable MAP were found in pasteurized milk [8] and in dried dairy products like powdered infant formula [9] and in raw fermented sausages [10], MAP are discussed as foodborne pathogens.

A similar pathology in the intestinal tissue of patients with intestinal tuberculosis and paratuberculosis was described more than a century ago. Recently, an association between MAP and Crohn’s disease was shown, initiating a discussion about a possible relationship of MAP in Crohn’s pathogenesis [11]. Johne’s and Crohn’s disease share clinical and histopathological similarities, MAP can survive standard pasteurization procedures and MAP antibodies can be detected in Crohn’s patients, where macrolide antibiotics ameliorate disease [12]. In contrast, genotypes of MAP isolated from cattle and man are different, there is a lack of evidence for uptake of contaminated food in respective patients and MAP cannot consistently be isolated from Crohn’s disease patients [13]. Neither in cattle, nor in humans MAP is transmitted from mother to child during pregnancy. But although MAP was detected widespread in many farms and different countries, the incidence of Johne’s disease in ruminants is marginal [14]. Bacteria can survive for 2–10 years without causing obvious symptoms of infection in cows [15]. As seen in cattle farms, susceptibility to MAP infection differs in human populations [16]. This points to a complex disease in which several pathogens, environmental factors and an inappropriate immune response in genetically susceptible hosts participate in the cause of disease [16]. Since an enhanced susceptibility of the host contributes to pathogenesis in other mycobacteria associated diseases (e.g. tuberculosis) [17], we wanted to gain further information about the interplay of MAP with the immune system in hosts with different immune capacities. This is also of interest because in cattle, the MAP eradication programmes that are solely based on hygiene management are not very successful [18]. This could indicate certain reservoir cows that host and spread MAP without developing any clinical signs.

Only little is known about the host-pathogen interaction of MAP and the immune system of its hosts [19]. Functional differences in these responses could lead to aberrant reactions in susceptible hosts [19].

Recently, we detected a functionally different immune capacity in 22% of cows from different herds in Germany using differential proteome analyses [20]. These immune deviant (ID) cows differ in their constitutive immune proteome and they regulate different master immune regulators upon polyclonal immune cell stimulation. The phenotype is functionally correlated with an increased prevalence of mastitis, indicating an impact on the ability to fight infections [20].

All living microorganisms are exposed to changing environmental parameters that define their habitats. Bacteria sense environmental changes and react to it with various stress response mechanisms [21]. To gain information about the pathogenic mechanisms of MAP and how they respond to different immune response signals of their hosts, we analysed the changes in the proteomes of MAP after co-incubation with primary peripheral blood derived leukocytes (PBL) of control and ID cows. Since the aim of this study was a better understanding of the host-pathogen response, we analysed the exoproteomes of MAP and the bovine peripheral blood derived lymphocytes.

The term exoproteome describes the protein content that can be found in the extracellular proximity of a given biological system [22]. These proteins arise from cellular secretion, other protein export mechanisms or cell lysis, but only the most stable proteins in this environment will remain abundant [22]. These proteins play roles in the organism’s survival in extreme habitats such as saline environments [23]. Investigating the exoproteome of the pathogen and the secretome of PBL provides expanded coverage of the repertoire of proteins secreted in the stress response of MAP to different immune responses and this is essential for understanding respective mechanisms. Accordingly, this study aimed at providing a better understanding of the interplay of *Mycobacterium avium* subsp. *paratuberculosis* (MAP) with the immune system on proteome level to identify the complex network of proteins involved in host-foodborne bacteria communication.

## 2. Materials and Methods

### 2.1. Mycobacterium avium subsp. paratuberculosis (MAP)

The bacterial strain used in these experiments was *Mycobacterium avium* subsp. *paratuberculosis* (DSM 44133), purchased from German Collection of Microorganisms and Cell Cultures (DSMZ, Braunschweig, Germany). MAP were grown on Herrold’s egg yolk agar (HEYM) (BD Biosciences, Heidelberg, Germany) for four weeks prior to harvesting for the co-incubation experiment. MAP were yielded through rinsing the agar of the cultivation tubes with phosphate buffered saline (PBS) and gentle scratching.

### 2.2. PBL isolation of control and immune deviant cows

Blood samples of cows were collected in tubes supplemented with 25.000 I.U. heparin. Blood was then diluted 1:2 with PBS pH 7.2 and subsequently layered on density gradient separating solution (Pancoll; PanBiotech, Aidenbach, Germany). After density gradient centrifugation (room temperature, 290 × g, 25 min, brake off), PBL were obtained from intermediate phase. Cells were then washed 3x in PBS (4°C). Withdrawal of blood was permitted by the local authority Regierung von Oberbayern, Munich, permit no. 55.2-1-54-2532.3-22-12. For determination of control or ID status, PBL were tested in *in vitro* proliferation assays as described [20]. Respective animals were tested at least 11 times, before being assigned to control or ID status. The animals were from a MAP-free farm and were tested negative for MAP antibodies and no MAP were detected in culture from feces.

### 2.3. Co-cultivation of MAP with primary bovine PBL

Primary bovine PBL (2×10^7^) of control and ID cows were cultivated in 6 well plates in 3 ml RPMI each. 15 µg of live MAP were added for 48 h to one well, control wells were not infected. After 48 h, three technical replicates per experiment were centrifuged for 10 min at 350 x g and the supernatants were filtered through 0,5 µm filters (= exoproteome) and stored at −20°C until filter aided sample preparation (FASP). Cell pellets were lysed and total protein content measured with Bradford assay.

### 2.4. Proteolysis and LC-MS/MS mass spectrometry

10µg total protein was digested with LysC and trypsin by filter-aided sample preparation (FASP) as described [24]. Acidified eluted peptides were analysed in the data-dependent mode on a Q Exactive HF mass spectrometer (Thermo Fisher Scientific, Bremen, Germany) online coupled to a UItimate 3000 RSLC nano-HPLC (Dionex). Samples were automatically injected and loaded onto the C18 trap column and after 5 min eluted and separated on the C18 analytical column (75µm IDx15cm, Acclaim PepMAP 100 C18. 100Å/size, LC Packings, Thermo Fisher Scientific, Bremen, Germany) by a 90min non-linear acetonitrile gradient at a flow rate of 250 nl/min. MS spectra were recorded at a resolution of 60000 and after each MS1 cycle, the 10 most abundant peptide ions were selected for fragmentation.

### 2.5. Protein identification and label-free quantification

Acquired MS spectra were imported into Progenesis software (version 2.5 Nonlinear Dynamics, Waters) and analyzed as previously described [6, 25, 26]. After alignment, peak picking, exclusion of features with charge state of 1 and > 7 and normalization, spectra were exported as Mascot Generic files and searched against a database containing all entries of *Mycobacterium avium* subspecies *paratuberculosis* from NCBI Protein database combined with the Ensembl bovine database (version 80) with Mascot (Matrix Science, Version 2.5.1). Search parameters used were 10 ppm peptide mass tolerance, 20 mmu fragment mass tolerance, one missed cleavage allowed, carbamidomethylation was set as fixed modification, and methionine oxidation and deamidation of asparagine and glutamine as variable modifications. Mascot integrated decoy database search was set to a false discovery rate (FDR) of 1% when searching was performed on the concatenated mgf files with a percolator ion score cut-off of 13 and an appropriate significance threshold p. Peptide assignment was reimported to Progenesis Software. All unique peptides allocated to a protein were considered for quantification. Proteins with a ratio of at least five-fold in normalized abundance between control and ID or samples were defined as differentially expressed.

### 2.6. Enriched pathway analyses

Abundances of the identified proteins were defined as differentially expressed based on the threshold of protein abundance ratio of ≥1.5 and their assignement to MAP or bovine exoproteome. Venn diagram was made with open source tool: http://bioinformatics.psb.ugent.be/webtools/Venn/. The protein–protein interaction network of differentially-accumulated proteins was analysed with GeneRanker of Genomatix Pathway System (GePS) software (version 3.10; Intrexon Bioinformatics GmbH, Munich, Germany; settings: Orthologous genes from H. sapiens were used for this ranking, species bovine, analysed were proteins with ≥ 5-fold change). Further bioinformatic analyses were conducted with open source software ShinyGO v0.50: http://ge-lab.org/go/ [27], P-value cutoff (FDR) was set to 0.05.

## 3. Results

### 3.1. Differentially abundant proteins in secretomes and exoproteomes of PBL from control and immune deviant hosts as response to in vitro infection with MAP

We investigated the immune response of primary PBL isolated from control and immune deviant animals. After 48 hours of in vitro infection with live MAP, we harvested the mixed secretomes and exoproteomes of PBL and MAP and analysed them with mass spectrometry. Overall, we identified 826 proteins (811 bovine and 15 MAP proteins). Cluster analysis confirmed significant differences in protein abundances between PBL of control and ID cow in host-pathogen response (Figure 1, Venn diagramm). In secretome of control cow, 90 proteins were differentially upregulated (Figure 1, ≥ 5-fold change of expression, blue circle) in contrast to 38 proteins that were higher abundant in ID cow (Figure 1, light red circle). In the MAP exoproteome, although small in overall numbers, different protein abundances were also detectable. In MAP co-incubated with control PBL, one protein (GroEl1) was found with protein abundance change and one protein (DnaK) was upregulated in exoproteome after co-incubation with ID PBL.

**Figure 1:**
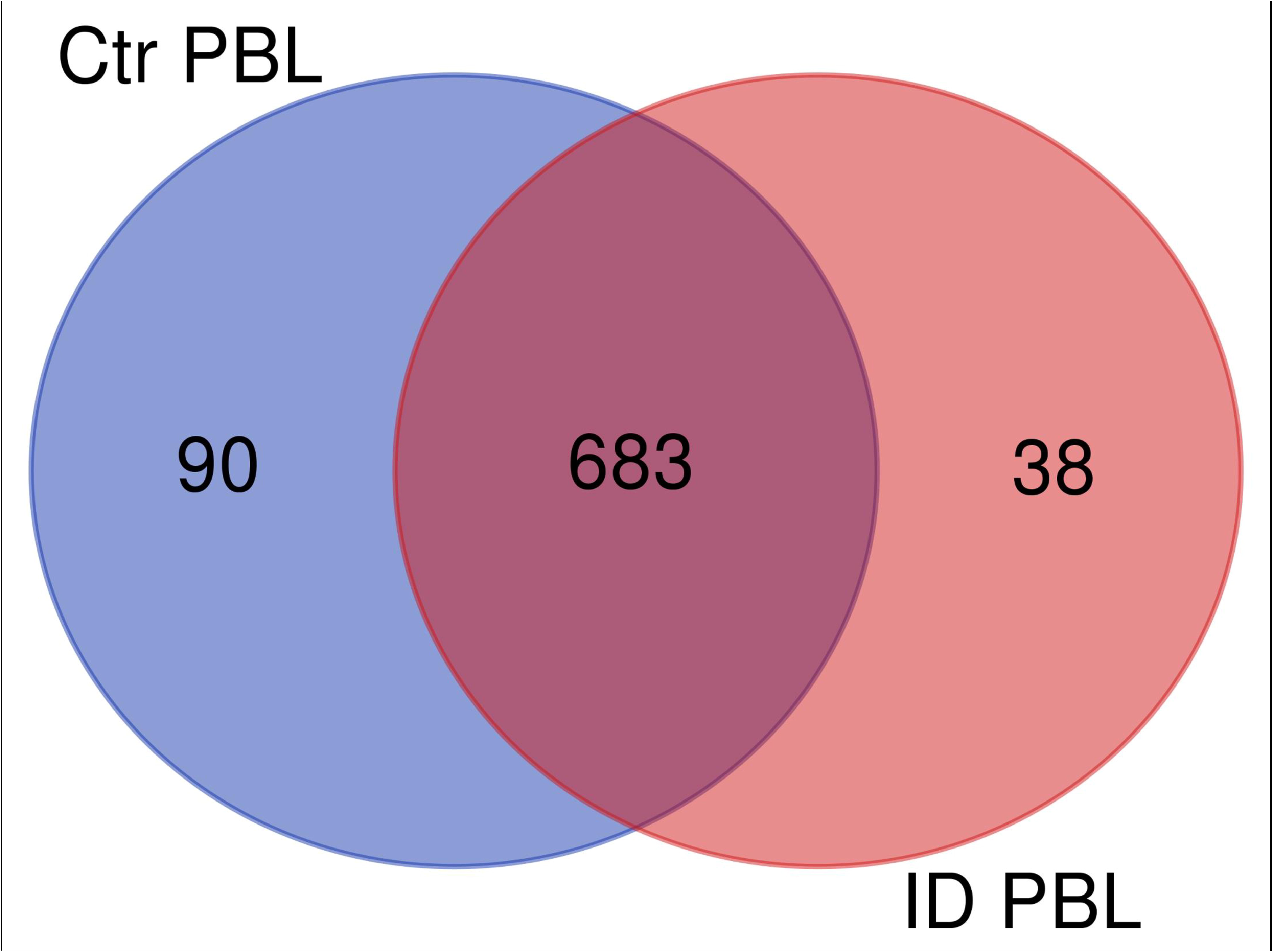
Overlap (Venn diagram) of differentially (≥ 5 fold) expressed proteins between secretomes/exoproteomes of control cow (blue) and ID cow (red). From a total of 811 identified proteins, 90 were higher abundant in control and 38 in ID.

### 3.2. Different biological pathways were regulated in the immune response in host PBL

Interpretation of large protein sets can be performed through enrichment analyses, using published information for examination of overrepresentation of a known set of genes within the input gene list [27, 28]. Since many gene ontology (GO) terms are related or redundant, we used a hierarchical clustering tree and network (ShinyGO v.050 [27]). The top regulated biological process pathways in the secretome/excretome of the control were all related to RNA splicing (Figure 2A). The top regulated immune pathways were cellular immune responses of various leukocyte subsets, chemotaxis and the reaction to IL-12 (Figure 2A).

**Figure 2:**
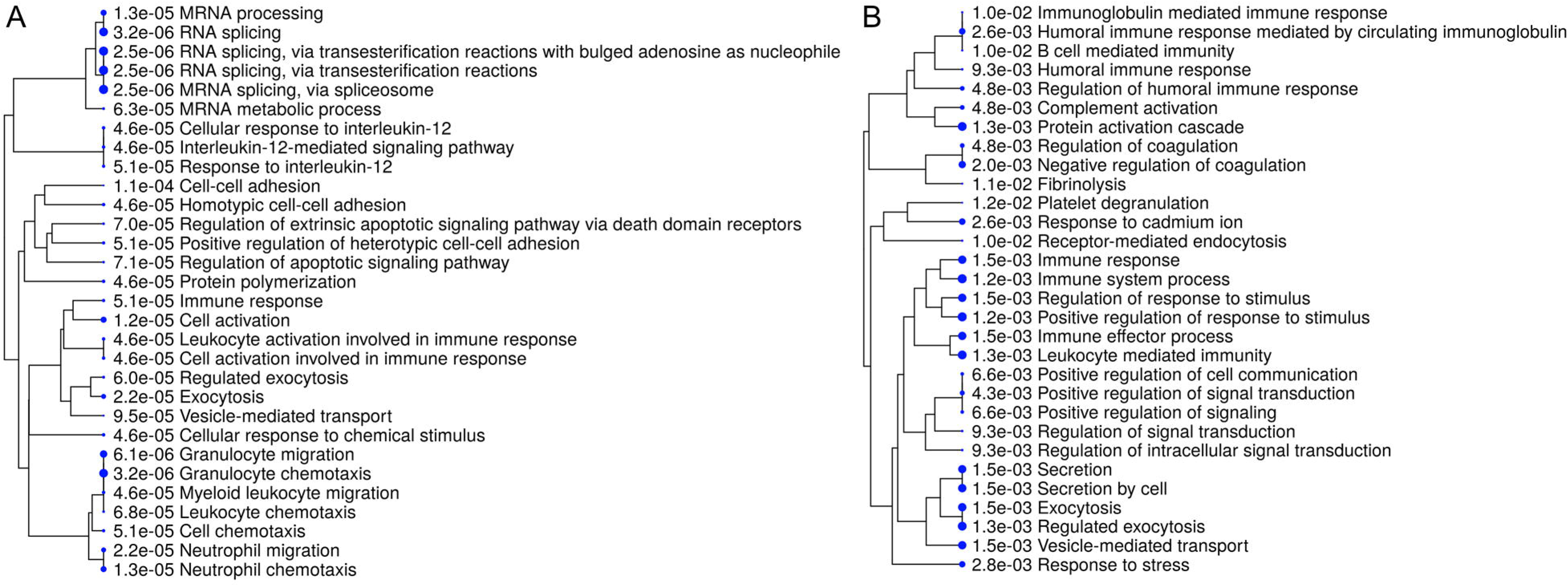
Hierarchical clustering tree and network of related GO terms of differentially expressed proteins in secretomes/exoproteomes of control PBL (A) and ID PBL (B) illustrates marked differences. GO terms are grouped together based on how many genes they share. The size of the solid circle corresponds to the enrichment false discovery rate.

In contrast, in secretome/excretome of ID PBL, the top regulated pathways were response to stress and immune system process (Figure 2B). The enriched immune pathways comprised distinct routes of immune response in ID PBL, focussing on closely related positive regulations of cellular functions, complement activation and humoral immune response (Figure 2B).

Visualization of network clearly indicates two major regulated networks in control PBL’s secretome (Figure 3A) and one distinct, major enriched network in ID proteins (Figure 3B).

**Figure 3:**
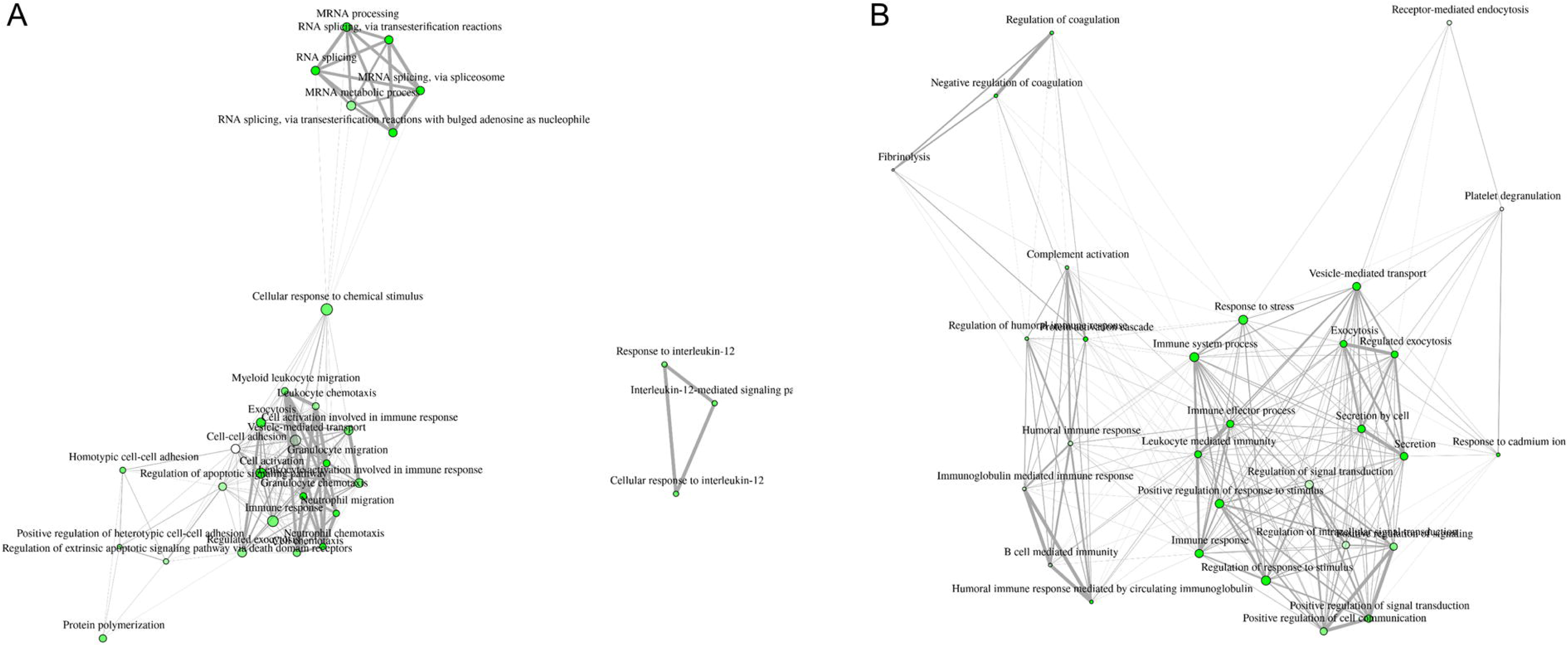
Visualization of overlapping relationships among enriched gene-sets revealed A) two major networks as shown by network view for enriched GO molecular component terms in control PBL after MAP infection and B) in ID PBL after MAP infection. Related GO terms are connected by a line whose thickness reflects percent of overlapping genes. Size of the nodes corresponds to number of genes.

### 3.3. GroEl1 and DnAK were differentially abundant in MAPs interacting with different hosts

Interestingly, there was also a clear difference in the regulated proteins of the pathogen itself, dependent on the co-cultivated host PBL (Table 1, supplementary table 2), although the low number of proteins was not sufficient for in depth enrichment analyses of the MAP exoproteomes. These differential proteome analyses clearly indicate substantial differences on protein level of the foodborne pathogen MAP in the interaction with different host immune capacities.

**Table 1:** Regulation of MAP proteins identified in secretomes/exoproteomes after co-incubation with control and ID PBL.

|  | Accession | Gene<br>symbol | Description | Ratio<br>MAP<br>Ctr/ID |
| --- | --- | --- | --- | --- |
| 1 | AAS06815 | GroEL1 | pep:novel<br>chromosome:GCA_000007865.1:Chromosome:473<br>3338:4734954:1 gene:MAP_4265<br>transcript:AAS06815 description: ""GroEL1"" | 6,07 |
| 2 | ETB04387 | F0F1 ATP<br>synthase<br>subunit<br>alpha | pep:novel<br>supercontig:GCA_000504785.1:contig000131:9486:<br>11150:1 gene:O979_07265 transcript:ETB04387<br>description: ""F0F1 ATP synthase subunit alpha"" | 3,26 |
| 3 | ETB05569 | DNA-bindin<br>g protein | pep:novel<br>supercontig:GCA_000504785.1:contig000072:8572:<br>9225:1 gene:O979_04085 transcript:ETB05569<br>description: ""DNA-binding protein"" | 2,47 |
| 4 | AAS06727 | RplN | pep:novel<br>chromosome:GCA_000007865.1:Chromosome:465<br>2017:4652385:1 gene:MAP_4177<br>transcript:AAS06727 description: ""RplN"" | 2,34 |
| 5 | AAS06486<br>;<br>ELP44387;<br>ETB08817 | GroEL2 | pep:novel<br>chromosome:GCA_000007865.1:Chromosome:439<br>5922:4397547:1 gene:MAP_3936<br>transcript:AAS06486 description: ""GroEL2"" | 1,12 |
| 6 | AAS06693 | Tuf | pep:novel<br>chromosome:GCA_000007865.1:Chromosome:462<br>0946:4622136:1 gene:MAP_4143<br>transcript:AAS06693 description: ""Tuf"" | 0,88 |
| 7 | ETB02420 | enoyl-CoA<br>hydratase | pep:novel<br>supercontig:GCA_000504785.1:contig000238:14619<br>:15410:1 gene:O979_11400 transcript:ETB02420<br>description: ""enoyl-CoA hydratase"" | 0,86 |
| 8 | ETB00930 | 2-isopropylm<br>alate<br>synthase | pep:novel<br>supercontig:GCA_000504845.1:contig000411:16560<br>:18359:-1 gene:O978_19245 transcript:ETB00930<br>description: ""2-isopropylmalate synthase"" | 0,75 |
| 9 | AAS06690 | RpsL | pep:novel<br>chromosome:GCA_000007865.1:Chromosome:461<br>7754:4618128:1 gene:MAP_4140<br>transcript:AAS06690 description: ""RpsL"" | 0,67 |
| 10 | AAS06681<br>;<br>ETB45918;<br>ETB45924 | RpoC | pep:novel<br>chromosome:GCA_000007865.1:Chromosome:460<br>6729:4610679:1 gene:MAP_4131<br>transcript:AAS06681 description: ""RpoC"" | 0,62 |
| 11 | AAS06680 | RpoB | pep:novel<br>chromosome:GCA_000007865.1:Chromosome:460<br>3087:4606683:1 gene:MAP_4130<br>transcript:AAS06680 description:""RpoB"" | 0,48 |
| 12 | AAS02778 | ClpC | pep:novel<br>chromosome:GCA_000007865.1:Chromosome:488<br>298:490829:1 gene:MAP_0461 transcript:AAS02778<br>description:""ClpC"" | 0,46 |
| 13 | ETB04840 | GTP-binding<br>protein YchF | pep:novel<br>supercontig:GCA_000504785.1:contig000104:1736:<br>2809:1 gene:O979_06015 transcript:ETB04840<br>description:""GTP-binding protein YchF"" | 0,43 |
| 14 | ETB04389<br>AAS04768 | ATP<br>synthase<br>subunit beta | pep:novel<br>supercontig:GCA_000504785.1:contig000131:12100<br>:13557:1 gene:O979_07275 transcript:ETB04389<br>description:""ATP synthase subunit beta"" | 0,26 |
| 15 | AAS06390 | DnaK | pep:novel<br>chromosome:GCA_000007865.1:Chromosome:429<br>5544:4297415:1 gene:MAP_3840<br>transcript:AAS06390 description:""DnaK"" | 0,04 |

## 4. Discussion

*Mycobacterium avium* subsp. *paratuberculosis* (MAP) could be a foodborne pathogen, because it is discussed in association with several diseases [29-31]. MAP were found in patients with inflammatory bowel diseases like Crohn’s disease and ulcerative colitis, as well as autoimmune diseases like type 1 diabetes, multiple sclerosis, rheumatoid arthritis and Hashimoto’s thyroiditis (Garvey, 2018), but so far, a causal association has not yet been proven in any of these cases. Since MAP can survive many of the standard techniques of food decontamination (e.g. pasteurization), they are regularly found alive in pasteurized milk (Gerrard et al., 2018) and in dried dairy products such as powdered infant formula (Botsaris et al., 2016). If MAP play a role in pathogenesis of respective diseases, we think that this must be associated with a certain type of susceptibility from these hosts. MAP is ubiquitously found and only a minor proportion of consumers (if at all) is affected by MAP-associated diseases. The same is true for cows: while they are often in contact with MAP, resulting in high frequencies of seropositive animals – approximately 20% and at least 3–5% in several countries [32, 33] – Johne’s disease incidence is very rare. For example, a total of 232 clinical cases of Johne’s disease were reported in Ireland from 1995 to 2002 [34], yielding an average annual rate of approximately 0.0005%, given a cattle population of six million [35]. A study examining environmental samples from 362 dairy farms located in all 10 provinces of Canada for detection of MAP by culture revealed true prevalence estimates of 66% for farms in Western Canada, 54% in Ontario, 24% in Québec, and 47% in Atlantic Canada [36].

From tuberculosis it is known that after infection with *Mycobacterium tuberculosis* (MTB), there is a 10% probability that the host will develop active tuberculosis and the bacterium may invade multiple organs [37]. Although 9 million new cases of active tuberculosis are still reported annually, an estimated one-third of the world is infected with MTB while remaining asymptomatic, defined as latent TB [38]. Among the individuals with latent TB, only 5–10% will develop active tuberculosis disease in course of their lifetime, because they effectively control the infection through their immune response [38]. This immune response after MTB infection is highly complex as the bacteria have intricate immune escape mechanisms [37]. For MAP, only little is known about host-pathogen reactions in general, and whether different immune responses exist, but a thorough examination of the respective mechanisms is of major importance to get functional data that allow a better understanding and a substantiated risk assessment. In the post-genomic era, proteomics represents a key discipline to perform in depth studies and identify the complex network of proteins involved in such host-bacteria communication. In this study, we analysed the exoproteomes/secretomes of MAP and host cow PBL with known, functionally different immune capacities [20] using differential proteome analyses after *in vitro* infection. Interestingly, there were significant differences in protein abundances secreted from control and ID PBL. Ninety proteins were ≥ 5 fold higher abundant in control PBL after interaction with MAP. In the control, the top regulated immune pathways described cell activation and chemotaxis of leukocytes as well as IL-12 mediated signalling pathways. For *M*. *paratuberculosis* it was shown, that IL-12 transcription is increased in infected bovine macrophages within 6 hours [39], probably to enhance the developing T cell response [40]. To our knowledge, the role of IL-12 in MAP infections of cows has not been analysed so far, but the IL-12 associated immune response should also be protective. In tuberculosis it was shown that the induction of protective IFN-γ T cell responses against primary *M*. *tuberculosis* infection clearly depends on IL-12 [41]. Mice lacking IL-12p40 cannot control the growth of the MTB bacterial infection [42]. These findings in mouse models were confirmed in man, where IL-12 was shown to be critical for preventing tuberculosis [43]. Therefore, we hypothesize that secretion of from IL-12indicates a protective immune response against MAP infection in control PBL. In ID PBL on the other hand, CCR4-NOT transcription complex, subunit 1 (CNOT1) was detected as highly abundant protein (supplemental table 1). This novel finding of CNOT1 regulation in the bovine immune system is very interesting, because CCR4-NOT complex members have recently been shown to function as regulators ensuring repression of the MHCII locus in human cell lines [44]. Poor MHC II expression can cause autoimmune or infectious diseases, since MHCII is indispensable for adequate immune responses [44]. Additionally, CNOT proteins also contribute to the downregulation of MHC class I gene expression by influencing transcription and mRNA degradation [45]. Enriched network analyses of secreted proteins from ID PBL revealed further major different functions of the 38 differentially abundant proteins (Figure 4B). Members of complement activation pathway were enriched in ID PBL. We think the regulated candidates in ID PBL merit further investigation in future studies to clarify whether they indicate an immune response in favour of MAP infection or just another way to successfully fight mycobacterial infections.

Interestingly, in the MAP exoproteome (although small in overall numbers) different protein abundances were also detectable. Exoproteomes arise from cellular secretion, other protein export mechanisms or cell lysis, but only the most stable proteins in this environment will remain abundant [22]. In MAP co-incubated with control PBL, GroEL1 showed 6-fold higher abundance. GroEL1 belongs to the family of 60 kDa heat shock proteins, also known as Cpn60s (GroELs) which are components of the essential protein folding machinery of the cell, but are also dominant antigens in many infectious diseases [46]. GroEL1 from MAP is highly immunogenic for cows [45]. The exact function of GroEL1 in MAP is not clarified so far, but it is highly similar to the respective protein in MTB (Rv3417c), where GroEL1 is important for bacterial survival under low aeration by affecting the expression of genes known for hypoxia response [46]. From co-cultivation of MAP with ID PBL, on the other hand, DnaK emerged as differentially abundant protein. DnaK is a HSP70 family chaperone protein with essential function in stress induced protein refolding and DnaK loss is accompanied by disruption of membrane structure and increased cell permeability [47]. DnaK is essentially required for cell growth in mycobacteria due to a lack of redundancy with other chaperone systems [47]. This finding from MAP – ID PBL co-cultivation points to regulation of important survival mechanisms in MAP. However, these results must be interpreted with care because our analysed MAP exoproteome only comprised 15 proteins.

Our data provide novel information about MAP-leukocyte interaction, adding to a more comprehensive picture of host-pathogen interactions. Co-incubation of MAP with cells from animals with different immune capacities led to significant differences in PBL secretomes and different immunological pathways enhanced in the hosts. In exoproteomes of respective MAPs, GroEL1 and DnaK were differentially abundant. These analyses gave a deeper insight into the different responses of host PBL and MAP bacteria. In this study, several novel proteins were identified with changed abundance in host-pathogen interaction. These candidates merit further investigations in the future to clarify their functional role in infection control.

## Supporting information

Supplemental Table 1_PBL

Supplemental Table 2_MAP

## Acknowledgments

The authors would like to thank Barbara Amann for excellent technical support.

## Author Contributions

C.D. conceived and designed the experiments; K.K., S.H., A.S. and C.H. performed the experiments; K.K., S.H., R.D., E.M. and C.D. analyzed the data; C.D. wrote the manuscript. All authors critically read the manuscript and approved the final version to be published.

## Abbreviations

CCR4: CCR4-NOT transcription complex, subunit 1
FASP: Filter-aided sample preparation
ID: Immune deviant
IL-12: Interleukin 12
HEYM: Herrold’s egg yolk agar
MAP: *Mycobacterium avium* subsp. *paratuberculosis*
MTB: *Mycobacterium tuberculosis*
PBL: Peripheral blood derived lymphocytes
PBS: Phosphate buffered saline

